# Quantitative Single-Cell Imaging of Efflux Pump Heterogeneity and Antibiotic Response Dynamics

**DOI:** 10.1101/2025.08.06.668869

**Authors:** Ruizhe Li, Somenath Bakshi

## Abstract

Understanding how bacterial cells survive antibiotic treatment requires tools capable of resolving dynamic physiological heterogeneity at single-cell resolution. Here, we present a high-throughput, lineage-resolved microfluidic imaging platform for quantifying the expression and spatial distribution of the AcrAB-TolC efflux pump in *Escherichia coli* during antibiotic exposure. The system is built around a custom-designed multilayer microfluidic device, fabricated using a direct-write photolithography protocol, which ensures uniform delivery of nutrients and antibiotics while confining thousands of individual cells in dead-end trenches. Brightfield imaging provides a label-free, high-temporal-resolution method for tracking cell growth and division, enabling accurate reconstruction of lineages over time. Efflux pump abundance is quantified using GFP-tagged AcrB, while a red fluorescent reporter driven by a constitutive promoter serves as a control to account for growth-dependent fluorescence artifacts. To analyse the sources of heterogeneity in efflux abundance and its consequences for antibiotic response dynamics, we developed a lineage-resolved analytical framework that compares cells of similar replicative age but differing spatial positions, and vice versa. This analysis was made possible by a novel machine-learning-based image processing pipeline, which enables robust cell segmentation, tracking, and lineage reconstruction across multiple fields of view. By extracting time-resolved single-cell data, this platform allows precise dissection of non-genetic variability in efflux activity and its role in determining survival outcomes, offering a powerful foundation for future quantitative studies of bacterial physiology under antibiotic stress.

## Introduction

Many bacterial species, particularly Gram-negative bacteria, tolerate toxic compounds through the action of energy-dependent, multicomponent efflux pumps that span both membranes and are powered by inner-membrane transporters (Du et al., 2018). Efflux pumps are especially central to bacterial survival under antibiotic stress, enabling cells to transiently reduce intracellular drug concentration (Poole, 2007). Yet, efflux activity is often heterogeneous within clonal populations, contributing to phenotypic variability in antibiotic susceptibility, recovery, and persistence (Sánchez-Romero and Casadesús, 2014). Resolving this heterogeneity, and its consequences, requires high-throughput, time-resolved measurements of both efflux levels and cellular responses at the single-cell level.

Several recent studies have illuminated the importance of efflux variability using single-cell platforms. For example, Bergmiller et al. (2017) used a microfluidic mother machine device to show that the multidrug efflux pump AcrAB-TolC partitions asymmetrically at cell division, conferring long-lived phenotypic memory that enhances antibiotic tolerance in aging mother cells. In parallel, El Meouche and Dunlop (2018) used agar pad–based time-lapse imaging and dual fluorescent reporters to uncover an inverse correlation between efflux expression and mismatch repair activity, suggesting that high-efflux cells may serve as transient mutators under stress. In this chapter, we present an experimental approach to revisit and extend the analysis of AcrAB-TolC efflux pump heterogeneity, using it as a case study to showcase the capabilities of our high-throughput, lineage-resolved single-cell imaging platform. By combining advanced multilayer microfluidics, time-resolved fluorescence and brightfield imaging, and machine learning-based lineage tracking, our approach offers new insights into the origins of efflux variability and its impact on antibiotic response and recovery. We begin by outlining the limitations of earlier methodologies, which motivates the design of our integrated experimental and analytical pipeline. Using the AcrAB-TolC system as a representative example, we demonstrate how this platform enables quantitative analysis of efflux pump abundance across lineages and reveals how lineage-dependent differences influence the antibiotic response of individual cells.

Historically, agar pad-based imaging platforms have been used to track individual bacteria over time (Stylianidou et al., 2016). While these are easy to implement, they suffer from spatial inhomogeneity in nutrient and drug delivery (Dusny et al., 2015). Local nutrient depletion and waste accumulation across the pad can introduce spatially structured growth differences, particularly problematic when using slow-maturing fluorescent reporters (Balleza et al., 2018). These fluorescent reporters can accumulate higher signal intensity in slow-growing cells—not due to increased promoter activity, but because slower growth allows more time for the fluorophores to mature before being diluted by cell division (Fig. 1a). For example, in the case of RFP variants with a maturation time of ∼40 minutes, only about 33% of the synthesized proteins are expected to be fluorescent in a cell doubling every 20 minutes. If a cell grows more slowly, with a doubling time of 40 minutes, this fraction increases to approximately 50%, making the cell appear brighter despite similar expression levels. Consequently, studies relying on such reporters in agar-based systems may inadvertently misattribute slow growth from nutrient heterogeneity as a sign of increased efflux pump abundance and associated burden. Furthermore, in microcolonies grown on agar pads, light bleedthrough between closely spaced cells, arising from diffraction during image formation, makes it difficult to accurately quantify physiological heterogeneity from fluorescence signals of gene-expression reporters (Hardo et al., 2024).

**Figure 1.**
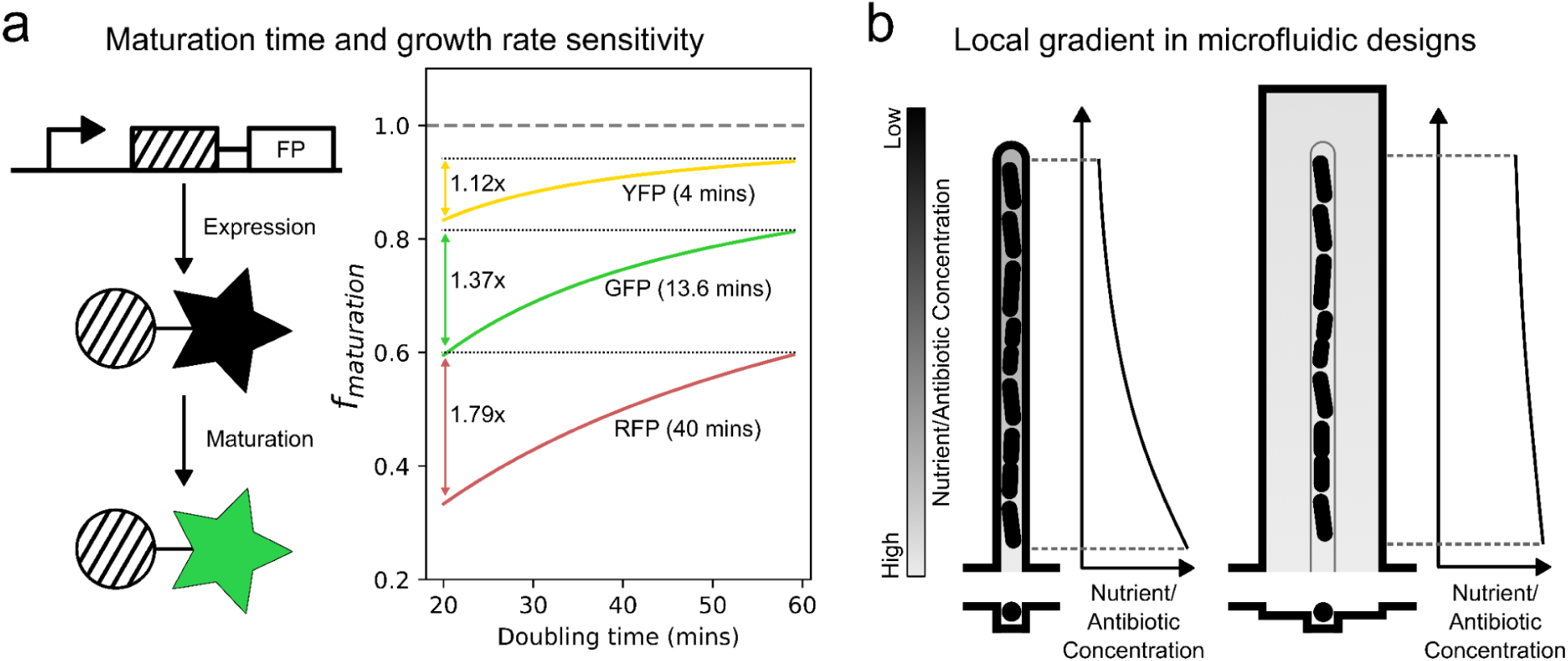
Design considerations for single-cell measurements of efflux pump expression and activity. *(a) Schematic of reporter protein expression and maturation, highlighting the impact of maturation kinetics on fluorescent signal interpretation. The plot (right) shows how maturation time interacts with cell growth rate to determine the steady-state fraction of mature fluorescent proteins. In fast-growing cells, dilution outpaces maturation, reducing signal intensity. For slow-maturing proteins such as RFP, this can result in artificially elevated fluorescence in slower-growing cells (e.g., ∼2× higher signal in 60 min vs. 20 min doubling time), potentially confounding interpretation of gene expression variability. In this study, we used a superfolder GFP variant (Bergmiller et al., 2017), which still exhibits growth-rate sensitivity. We recommend the use of faster-maturing fluorophores, such as mVenus NB, which matures in under 4 minutes and is more robust to growth rate differences. Maturation data were taken from the Balleza et al. (2018)*. *(b) Schematic illustrating limitations of the standard mother machine design. Nutrients and antibiotics enter the cell trench diffusively through the open end, but their concentrations decline along the trench due to cellular uptake and binding, particularly in the presence of a stack of dividing cells beneath the mother cell. This results in unpredictable and spatially varying physiological conditions at the closed end, complicating single-cell measurements. This effect can be mitigated by introducing adjacent shallow side trenches (right), which enhance lateral diffusion and improve chemical delivery to the mother cell*.

Microfluidic devices, particularly mother machine architectures, have since become the standard for single-cell studies of bacterial dynamics (Wang et al., 2010). These devices allow for precise environmental control, with laminar flow delivering well-defined media conditions uniformly across thousands of trenches. This enables robust experimental perturbations, such as transitions from media without antibiotics to media containing antibiotics and back, allowing systematic comparison of cellular responses and recovery kinetics. When properly designed, these platforms provide the statistical power and reproducibility required to quantify heterogeneity in antibiotic response across lineages. Additionally, it has been recently demonstrated that the inter-trench spacing in the mother machine design can be adjusted to alleviate the light bleedthrough effects discussed above, ensuring reliable quantification of the underlying phenotypic heterogeneity (Bakshi et al., 2021; Hardo et al., 2024).

Even in mother machine-type microfluidic devices, designed to provide uniform media access from the main flow channel, growth and treatment conditions within a trench can remain non-uniform. The mother cell, located at the closed end of the trench, is separated from the flow channel by a column of daughter cells. This arrangement can lead to local gradients in drug exposure due to differences in cell packing density, permeability, or metabolic activity (Fig. 1b). As a result, even when treatment is consistent across trenches, cell-to-cell variability within a trench may arise from subtle environmental inhomogeneities. Previously, such artefacts with the regular mother machine design were identified in nutrient profile and resultant cell resuscitation dynamics (Bakshi et al., 2021) and in oxidative stress response in bacteria (Choudhary et al., 2023). Gradients of nutrients and antibiotics along the trench can confound interpretation of single-cell responses. For instance, cells located near the closed end of the trench, typically those with older poles, may experience lower antibiotic concentrations due to diffusion limitations, giving the false impression of enhanced protection that is unrelated to their physiological state or replicative age.

To disentangle true biological heterogeneity from such spatial artifacts, we advocate a lineage-resolved analysis. By simultaneously tracking both the spatial position and lineage identity of cells, it becomes possible to compare cell pairs that share similar expression levels or replicative age but differ in position, or vice versa. This approach enables clear attribution of phenotypic differences, such as antibiotic survival or efflux pump expression, to either intrinsic factors like pole age or extrinsic factors such as differential access to treatment. For example, a cell deeper in the trench may receive a lower antibiotic dose than one near the channel opening, but by using lineage data to track pole age and efflux pump inheritance, one can determine whether enhanced survival arises from inherited protein levels or simply reduced antibiotic exposure. Nonetheless, it remains advisable to use improved microfluidic designs, such as the mother machine with side trench design proposed by Norman et al. (2013), that minimize environmental gradients from the outset (Fig. 1b). In this chapter, we rely on an optimised version of this design to mitigate the effects of antibiotic gradients, and combine lineage tracking, to precisely quantify the benefits of accumulated pumps in cells with older poles.

A further challenge in high-resolution time-lapse studies lies in balancing temporal resolution and phototoxicity. Frequent imaging of fluorescent reporters is often limited by bleaching and photodamage. Moreover, expressing reporters imposes a metabolic burden, which itself introduces heterogeneity in growth and stress response. To address this, here we developed a pipeline that uses brightfield-based segmentation and tracking, enabled by a machine learning algorithm trained on synthetic data generated from trench-like microfluidic environments (Hardo et al., 2022). This allows us to obtain high time-resolution imaging (1-2 min/frame) without relying on frequent fluorescence acquisition. Fluorescent reporters can then be imaged less frequently with low exposure and binned images, enabling accurate measurements of efflux pump levels with minimal perturbation. A predictive lineage reconstruction algorithm can then be used to link cell identities across frames, ensuring reliable tracking even under occasional segmentation uncertainty (Wedd et al., 2024). The proposed and developed pipeline for processing brightfield images also makes the method widely applicable to natural isolates and knockouts, obviating the need for cloning fluorescent segmentation markers in those cells.

### Chapter Overview

This chapter presents an end-to-end methodology for performing time-resolved, high-throughput, single-cell analysis of efflux pump expression and antibiotic response using microfluidic devices. We focus on the AcrAB-TolC pump of *E. coli*, which is a model efflux pump for gram-negative organisms (Du et al., 2014), using the reporter systems developed by the work of Bergmiller et al. (Bergmiller et al., 2017), to revisit the analysis of the origin and consequence of the heterogeneous abundance of efflux pumps across cells in the presence and absence of antibiotics. We employ advanced design of the microfluidic device for robust control of the growth and treatment environment, perform high throughput multichannel imaging studies to track growth, death, age, morphology, and gene expression dynamics of individual cells, and develop new methodologies for identifying and tracking individual cells and their phenotypic properties in a lineage-resolved manner with high temporal resolution. We cover each step of the process, from experimental design to quantitative analysis:

- Device Fabrication: Design and fabrication of mother machine-type microfluidic chips using photolithography and soft lithography methods.
- Sample Preparation and Cell Loading: Preparation of bacterial cultures and protocols for controlled loading into microfluidic trenches.
- Experimental Setup and Data Acquisition: Configuration of tubing, reservoirs, and valves to enable precise switching from media without antibiotics to antibiotic-containing media and back. Setup for time-lapse acquisition with optimized timing and exposure settings.
- Data Storage and Compression: Strategies for efficient data management of long time-lapse movies, including cropping and trench-wise compression.
- Image Processing: Segmentation and Feature Extraction: Machine-learning–based segmentation pipeline using synthetic training data to extract cell masks from brightfield images. Quantification of cell size, growth rate, division time, and efflux reporter fluorescence.
- Lineage Tracking and Reconstruction: Predictive algorithms to assign cell identities across frames and reconstruct lineages through growth and division events.

Throughout this chapter, we emphasize strategies to ensure that microfluidic single-cell studies of efflux pump heterogeneity are accurate, interpretable, and reproducible. These include:

- Microfluidic designs that minimize or control for intra-trench diffusion effects.
- Use of fast-maturing, low-background reporters, ideally in ratiometric or dual-color configurations.
- Analytical frameworks that integrate lineage tracking and spatial context into the quantification of fluorescence, growth, and drug response.

By grounding experimental design in an awareness of physical and biological confounders, researchers can obtain more faithful measurements of how efflux heterogeneity across the cell population shapes bacterial adaptation and drug response. By the end of this chapter, readers will have a complete experimental and computational framework to perform rigorous, artefact-resistant single-cell analysis of efflux pump heterogeneity, enabling insights into how molecular mechanisms of resistance translate into cellular behaviors under antibiotic stress.

### Microfabrication of Multilayer Microfluidic Devices for Single-Cell Antibiotic Treatment Assays

In this chapter, we describe the complete microfabrication workflow used to construct the microfluidic devices for single-cell efflux pump assays. These devices are fabricated in two main stages:

1. Hard lithography to produce a multilayer SU-8 master mold on a silicon wafer;
2. Soft lithography to replicate the master in PDMS (polydimethylsiloxane) and assemble it into a functional microfluidic chip.

The resulting devices enable precise control over bacterial microenvironments and support high-resolution imaging of individual cells over extended time periods under dynamic media conditions. The same fabrication pipeline is used for additional modules such as gradient generators.

The devices consist of three distinct layers:

1. A 0.5 µm-thick side trench layer, which surrounds the cell-trapping region and facilitates efficient media and antibiotic diffusion;
2. A 0.8 µm-thick cell trench layer, designed to capture and constrain rod-shaped bacterial cells in a monolayer;
3. A 25 µm-thick flow layer, which delivers nutrients and antibiotics across the device.

This modular layering enables better control over the physical microenvironment of each cell, while maintaining high optical compatibility and flow stability. The width and depth of the side trench layer can be adjusted to improve diffusive access of nutrients and antibiotics deeper into the trenches (see Figure 2 for details). The main cell trench layer can therefore be kept narrow and shallow, which reduces the movement of cells in the horizontal (which could complicate inter-channel image registration) and vertical directions (which could affect Z focus and image sharpness). These three layers are constructed sequentially on a silicon wafer using SU-8 photoresist formulations with different viscosity and spin parameters. The final SU-8 structure serves as a master mold for PDMS casting.

**Figure 2.**
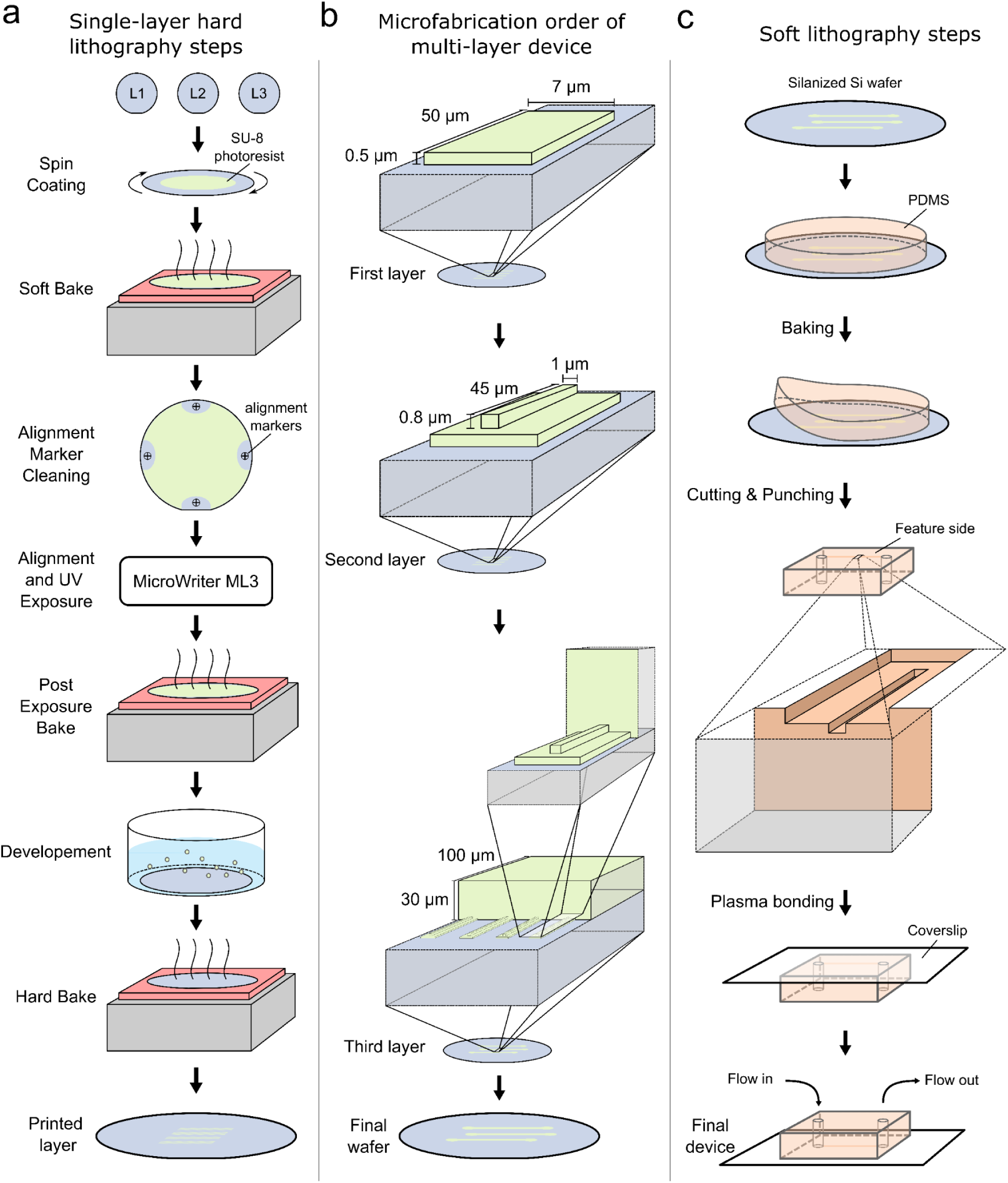
Hard and soft lithography of multilayer microfluidic devices. *(a)* Left: Schematic of the multilayer photolithography process used to fabricate the silicon master mold. The device used in this study, a mother machine variant with side trenches, requires three sequentially aligned SU-8 layers: L1 for the shallow side trenches, L2 for the cell trenches, and L3 for the main flow lane. Each layer is patterned using a photoresist of appropriate thickness (see Table 1) and aligned using dedicated registration markers. All layers were printed using a MicroWriter ML3 Pro direct-write photolithography system. *(b)* Middle: Overview of the hard lithography sequence, showing the order of layer printing and key dimensions for each feature. The final silicon wafer contains multiple device patterns, each comprising several flow lanes with independent inlet and outlet ports. Each flow lane houses over 1,000 cell trenches flanked by shallow side trenches that facilitate nutrient and treatment delivery. *(c)* Right: Soft lithography process for casting PDMS from the patterned master. The cured PDMS is cut into individual chips, and inlets/outlets for each flow lane are punched prior to plasma bonding with a glass coverslip of suitable thickness. The final bonded device supports multiple experiments in parallel, each conducted in an isolated flow lane.

**Table 1:** SU-8 photoresist selection and spin parameters for multilayer device fabrication.

| Layer | Target Thickness | SU-8 Type | Spin Programme (rpm/acceleration/time) | Soft Bake | Exposure Settings | Post-Exposure Bake | Comments |
| --- | --- | --- | --- | --- | --- | --- | --- |
| Alignment markers | ~1.0 $\mu\text{m}$ | SU-8 2001 | 500/100/10 + 1000/300/60 | 1 min @ 65 °C + 4 min @ 95 °C + 1 min @ 65 °C | 1350 mJ/cm<sup>2</sup> at -2 $\mu\text{m}$ focus | 1 min @ 65 °C + 6 min @ 95 °C + 1 min @ 65 °C | Expose first to enable alignment of subsequent layers |
| Side trench layer | ~0.5 $\mu\text{m}$ | SU-8 2000.5 | 500/100/5 + 2300/300/45 | 1 min @ 65 °C + 1 min @ 95 °C + 1 min @ 65 °C | 3500 mJ/cm<sup>2</sup> at -2 $\mu\text{m}$ focus | 1 min @ 65 °C + 5 min @ 95 °C + 1 min @ 65 °C | Requires high-resolution exposure and accurate alignment. |
| Cell trench layer | ~1.0 $\mu\text{m}$ | SU-8 2001 | 500/100/10 + 1000/300/60 (or 2000/300/60 for 0.8 $\mu\text{m}$) | 1 min @ 65 °C + 4 min @ 95 °C + 1 min @ 65 °C | 1250 mJ/cm<sup>2</sup> at -2 $\mu\text{m}$ focus | 1 min @ 65 °C + 10 min @ 95 °C + 1 min @ 65 °C | Can be combined with side trench with higher rpm to reduce thickness. |
| Flow layer | ~25 $\mu\text{m}$ | SU-8 2025 (or SU-8 2075 for >50 $\mu\text{m}$) | 500/100/10 + 4000/300/30 | 1 min @ 65 °C + 5 min @ 95 °C | 150 mJ/cm<sup>2</sup> at -1 $\mu\text{m}$ focus | 1 min @ 65 °C + 6 min @ 95 °C | Large features; use the low-magnification objective for exposure. Requires slightly stronger agitation during development. |

### Basic Steps in Hard Lithography: Fabrication of the SU-8 Master Mold

The fabrication process involves sequential patterning of SU-8 negative photoresist layers onto a clean silicon wafer, with careful alignment between each layer. The protocol is outlined below and illustrated schematically in Figure 2 a,b.

***Step-by-Step Protocol***:

a. Spin Coating: A uniform layer of SU-8 photoresist is applied to the silicon wafer using a spin coater. The specific SU-8 formulation and spin parameters depend on the target layer thickness (e.g., SU-8 2000.5 for 0.5 µm, SU-8 2001 for 1 µm, SU-8 2025 for 25 µm). Detailed specifications are provided in table 1.
b. Soft Bake: The coated wafer is baked on a hot plate to evaporate solvents and partially cure the resist. The soft bake time is kept consistent with the photoresist’s processing guidelines, as it is intended to evaporate the solvent, which depends on the chemical composition of the specific SU-8 photoresist for each layer.
c. Alignment Marker Cleaning: Before aligning subsequent layers, the area around the alignment marker is selectively cleaned using SU-8 developer to improve contrast and visibility during alignment.
d. Alignment and UV Exposure: UV patterning was carried out using the MicroWriter ML3 Pro system (Durham Magneto Optics), a high-resolution, maskless lithography tool. This system uses a digital micromirror device (DMD) to project 365 nm UV light in precisely stitched write fields. Alignment and focusing are guided by a computer-vision–assisted microscope objective. In this work, a 20× objective was used, providing approximately 0.6 µm lateral resolution in the xy plane. The MicroWriter’s computer-vision system accurately detects alignment features on the wafer by locating the centers of four predefined markers. A projective transformation is then applied to align and map the write fields onto the wafer surface, ensuring precise layer registration during multi-layer photolithography.
e. Post-Exposure Bake: A secondary bake is performed to complete crosslinking of the exposed photoresist.
f. Development: The wafer is immersed in SU-8 developer to remove unexposed regions of the photoresist.
g. Rinse and Dry: Developed wafers are rinsed with isopropanol (IPA) and dried with filtered nitrogen or air.
h. Hard Bake: A final hard bake is used to enhance the mechanical robustness and chemical resistance of the master.

### Multilayer Fabrication Workflow

These basic steps are repeated sequentially for each photoresist layer. The order of layer construction is critical to preserve alignment and structural integrity:

Step 1: Expose and bake alignment markers directly onto the wafer to serve as fixed reference points.

Step 2: Pattern the side trench layer (0.5 µm) using SU-8 2000.5 and a high-resolution objective. Step 3: Pattern the cell trench layer (1 µm) using SU-8 2001 with precise alignment to the previous layer.

Step 4: Pattern the main flow channel layer (25 µm) using SU-8 2025 or SU-8 2075, with a low-magnification objective for larger features.

Between each layer, careful cleaning, visual alignment using the MicroWriter’s overlay system, and verification of overlay accuracy are performed to ensure feature integrity. Misalignment across layers can severely impact trench geometry and flow dynamics, and therefore all designs are calibrated to minimize stitching and registration errors.

To achieve the desired geometry of the microfluidics features for our application, the correct combination of settings in the protocol must be optimised. Precise control over the thickness of the exposed photoresist and the shape of the features is essential for culturing bacteria in an organised manner. The thickness of the resulting features is measured using a profilometer and the shape is initially inspected under a microscope and later can be accurately measured using Atomic Force Microscopy (AFM). Additionally, the bonding between the photoresist and the wafer needs to be optimised. This bonding performance is highly dependent on the design of the features to be exposed. A well-optimised combination of settings should ensure that the bonding is sufficient without overexposing or underexposing the features. In some cases, features may need to be slightly overexposed to ensure good bonding to the substrate. If this is the case, the design can be deliberately adjusted to be smaller or modified to further improve the contact surface for bonding. Additionally, it is possible to improve bonding by performing an extra plasma treatment step before spin coating.

### Soft Lithography: Replicating the Master Mold and Device Assembly

Once the SU-8 master is completed, PDMS replicas of the microfluidic device are cast and bonded to glass substrates to create functional chips. Before the first PDMS casting, the silicon wafer with the microfluidic features needs to be secured onto an aluminium disc of identical diameter using heat-resistant epoxy to provide mechanical support. The silicon wafer then undergoes silanization to protect the SU-8 photoresists.

*Silanization Protocol (adapted from Microfabrication Core Facility, Harvard Medical School)*:

a. Clean the wafer with compressed air to remove dust particles. The wafer can be cleaned further by sequential washes of acetone, isopropanol (IPA), and distilled water, followed by drying with compressed air.
b. Place the wafer in a fume hood, next to an aluminum foil cap containing two drops of silanizing agent (perfluorooctyltrichlorosilane, FDTS).
c. Cover both the wafer and aluminium cap with a desiccator lid labeled “silanization” and leave at room temperature for 30 minutes to form a self assembled monolayer. This coating reduces surface energy and prevents sticking of the PDMS.
d. After silanization, transfer only the wafer onto a 150⁰C hotplate in the fume hood for 10 minutes to cure and evaporate the residual silanizing agents.

### PDMS Replica Molding

Polydimethylsiloxane (PDMS) replicas were prepared using Sylgard 184 elastomer kit (Dow Corning) at a base-to-curing agent weight ratio of 10:1. For a 2-inch silicon wafer, 10 g of base and 1 g of curing agent were mixed thoroughly in a clean disposable cup using a spatula. The mixture was then degassed in a desiccator under vacuum for approximately 30 minutes to eliminate air bubbles, carefully monitoring to avoid overflow by intermittently closing the vacuum valve if necessary. Meanwhile, a folded aluminum foil (2-4 layers) was wrapped around the wafer to create a no-leak container, exposing only the patterned surface. The degassed PDMS was poured onto the wafer from close range to minimize bubble formation and spread evenly to form a layer ∼0.5 cm thick. A second degassing step was performed with the foil-wrapped wafer to remove residual bubbles. The PDMS was then cured by baking at 95 °C for 1 hour. After cooling, the foil was carefully removed by cutting an “X” on the back side of the wafer and peeling the aluminum away. The cured PDMS layer was gently peeled off, avoiding fractures near microfeatures, and individual chips were excised on a cutting mat. Inlet and outlet holes were punched from the feature side using a 0.75 mm ID biopsy punch under a stereomicroscope. Debris was cleared from the punch between each use to prevent cracking.

### Cleaning of PDMS Replicas and Coverslips

Debris around the inlets and outlets was removed using Scotch tape. PDMS replicas were cleaned by sonicating in isopropanol for 30 minutes in a pre-cleaned beaker (washed with ethanol, dried with lint-free wipes, and compressed air), with the feature side facing up and the beaker covered with aluminum foil. The features were then dried with compressed air, taking care as the surface becomes slippery. Replicas were rinsed with Milli-Q water, followed by a second 30-minute sonication in Milli-Q water. For floating PDMS (due to hydrophobicity), replicas were placed feature side down to ensure contact with water. After cleaning, replicas were dried using an air gun and baked at 65 °C for 1 hour or 95 °C for 30 minutes to remove residual moisture.

Glass coverslips (22 mm × 50 mm, thickness 0.13–0.17 mm) were sonicated in 1 M KOH for 20 minutes using a coverslip holder in a clean beaker. Coverslips were then rinsed multiple times with Milli-Q water to remove all residual KOH and prevent crystallization upon drying. A second sonication in Milli-Q water was performed for another 20 minutes. After rinsing, coverslips were dried individually by blowing tangentially with compressed air and baked at 65 °C for 1 hour or 95 °C for 30 minutes.

### Bonding of PDMS Chips to Glass Coverslips

Bonding was performed using oxygen plasma treatment (Low-pressure plasma systems, model Diener Zepto, Diener Electronics). PDMS replicas (feature side up) and prepared coverslips were placed on an acrylic sheet inside the vacuum chamber of the plasma system. The plasma system was then turned on and allowed to initialize for ∼2 minutes with all valves closed. After closing the chamber, the vacuum pump was activated, and once a vacuum of 0.1 mbar was reached, the air valve was slowly opened to raise the pressure to 0.7 mbar. Plasma was applied at 70 W (35% of 200 W capacity) for 2 minutes while monitoring the magenta glow indicating plasma generation.

After treatment, the plasma and air valves were turned off, ventilation was opened to return to atmospheric pressure, and the chamber was opened immediately after that. Bonding was achieved by bringing the feature side of the PDMS mold into direct contact with the glass coverslip surface. Successful bonding was confirmed by the rapid propagation of the contact front, visible as a darkening of the interface due to the exclusion of air between the surfaces. The assembled chips were placed on a 95 °C hotplate for 5 minutes to enhance bonding and then baked for an additional hour at 95 °C to restore PDMS hydrophobicity.

### Sample Preparation and Cell Loading

With the microfluidic device fabricated and bonded to a glass substrate as described in the previous chapter, the next step is to prepare and load bacterial cultures into the cell trench region of the device. To ensure smooth, clog-free loading into the sub-micron trenches, we take several measures that depart from conventional approaches:

Cultures are grown overnight in the presence of a mild non-ionic surfactant, Pluronic F-108, which prevents cell aggregation and promotes uniform single-cell suspensions. This is essential for reliable entry of rod-shaped bacterial cells into narrow trench geometries.

Prior to loading, overnight bacterial cultures are concentrated 10× and introduced into the microfluidic device via gel-loading pipette tips inserted into the inlet and outlet ports. Once the cell suspension is pipetted into the flow channel, the tips are trimmed to a short length and sealed by gently melting the cut ends with a heated spatula. This sealing step prevents evaporation during the loading period and stabilizes conditions inside the device.

Unlike traditional mother machine devices, which typically require centrifugation to force dense cultures into dead-end trenches (Thiermann et al., 2024), our multilayer side-trench design enables convection-assisted loading. This approach eliminates the risk of coverglass cracking, a common failure during centrifuge-based loading, and allows cells to enter the trenches gradually and gently. As the concentrated culture flows into the main channel, it displaces air from the trench layer. Capillary action, enhanced by slow convection from evaporation through the porous PDMS, promotes continuous cell entry into the narrow cell trenches over time.

To minimize the introduction of air bubbles, which can disrupt loading and lead to evaporation-induced osmotic shifts in the trenches, it is critical to fully seal the flow channel after introducing cells. Bubbles can trap liquid in trenches and cause solute concentrations to rise as water evaporates, leading to osmotic stress. Cutting and simultaneously melting the gel-loading tips with a hot spatula ensures a secure, bubble-free seal at both inlet and outlet ends.

### Experimental Setup and Data Acquisition

After loading cells into the dead-end trenches of the microfluidic device, a flow path is established using a syringe pre-filled with the desired growth medium supplemented with 0.08% Pluronic F-108 (to prevent cell adhesion). Previously, we have verified that this concentration of Pluronic F-108 does not impact cell growth (Bakshi et al., 2021). The syringe is connected to high-strength silicone tubing (inner diameter: 0.50 mm; wall thickness: 0.50 mm) via a 21-gauge blunt needle. For interfacing the tubing with the microfluidic device, two 21-gauge blunt needles bent to a smooth 90° angle are used to ensure low-stress connections at the inlet and outlet ports.

Prior to connecting the flow path to the device, the entire tubing and needle system is sterilized by sequential flushing with 20% bleach, 20% ethanol, and distilled water. After this cleaning step, the flow path is flushed with the experimental medium to eliminate air bubbles from the tubing and needle connections. All effluent is discarded into a waste container containing 70% ethanol to ensure biosafety. Once the system is bubble-free and clean, the syringe is mounted on a syringe pump set to deliver medium at a flow rate of 15 μL/min.

To connect the device, gel-loading pipette tips (used earlier for cell loading) are carefully removed on a flat surface. During this step, one finger is gently pressed against the PDMS near each tip to prevent bending or detaching the glass coverslip. The outlet connection is made first by gently inserting the bent needle into the outlet hole. For the inlet, the pump is turned on before inserting the needle to ensure a smooth flow and prevent air entry. The connected device is then fixed onto the microscope stage using adhesive tape, and aligned with the principal axis of the camera by rotating the circular stage insert.

Once the assembly is complete, the Nikon Ti2 microscope incubator is closed to allow the system to reach thermal equilibrium. Stage positions corresponding to individual fields of view (FOVs) are then selected and recorded using Nikon’s NIS-Elements software, which enables automated multi-position, multi-color, time-lapse acquisition. For each FOV, a reference Z-position is saved to activate Nikon’s Perfect Focus System (PFS), which maintains consistent focus throughout the experiment.

The image acquisition parameters, including number of FOVs, time points per hour, and duration, are defined in the software. At each time point, images are acquired in multiple channels: brightfield (for cell segmentation), green fluorescence (to monitor GFP-tagged AcrB efflux pump), and red fluorescence (to track mCherry expressed from the control promoter pR, derived from the λ-phage right promoter) (Bergmiller et al., 2017). All image data is saved as a single multi-dimensional ND2 file.

The entire experiment, including pre-treatment growth, antibiotic exposure, and post-treatment recovery, is recorded in a continuous time-lapse within the same file. Media switching events are performed manually and their time points are logged for later alignment with image frames during analysis. Following the pre-treatment growth phase, the medium is switched to one containing a sub-MIC concentration of the translation inhibitor chloramphenicol (4 μg/mL) for 17 hours to induce stress (MIC 12 ug/mL). After the treatment phase, the system is flushed with antibiotic-free medium to capture recovery dynamics over time.

### Data Storage and Compression

Raw time-lapse movie captured by the Nikon imaging system is stored as a ND2 file, which contains multi-dimensional image data and metadata in one place. The initial data compression step involves extracting these into individual field of view (FOV) images for each time point and imaging channel using a custom Python script (Wedd, 2023) with the nd2 package (https://pypi.org/project/nd2/). Extracted images are organized into a directory with consistent naming conventions (field of view number + imaging channel + frame number, all automatically detected from the metadata). This extraction step can be parallelized in the script for efficient processing of large experimental datasets. Metadata including relevant image acquisition settings is also extracted to ensure no loss of experimental details. Extracted images are then corrected for rotation and registered for spatial drift, addressing temporal misalignments caused by mechanical stage movement or mismatched thermal expansion of the chip and the objective (Hardo, 2025a). The schematic of this pipeline is shown in the Fig. 3a and more details can be found in Hardo (2025b), Hardo et al. (2022).

**Figure 3.**
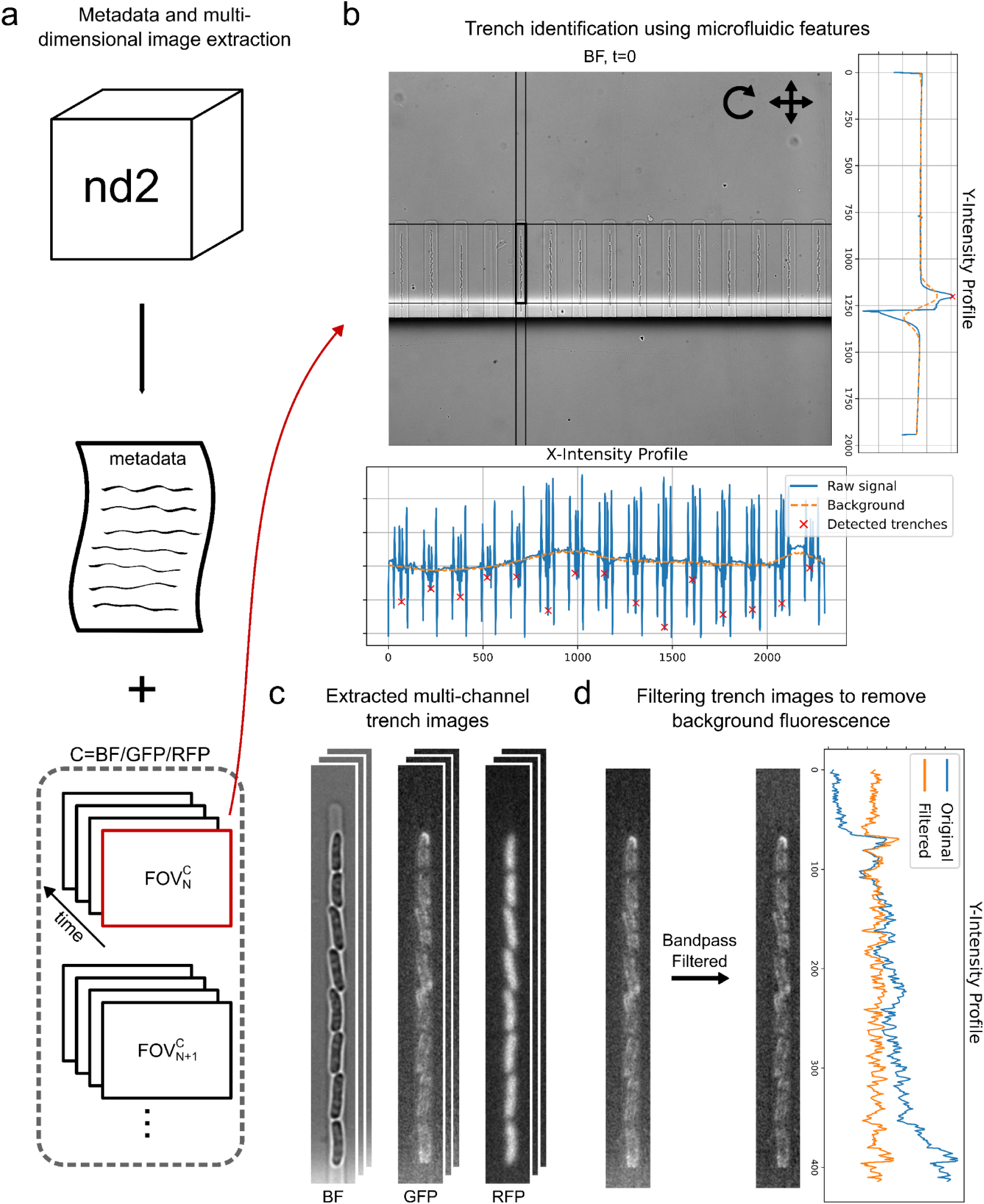
Data compression and preprocessing for time-resolved single-cell analysis. *(a) Schematic illustrating the extraction of metadata and multi-dimensional images from raw ND2 files. Images extracted correspond to specific fields of view (FOV), imaging channels, and time frames. This step prepares the images for preprocessing, while preserving all relevant metadata*. *(b) Key preprocessing steps to isolate trench images. FOV images are first rotated and registered to maintain consistent alignment of trenches over time. Mean X and Y intensity profiles from the bright field channel (averaged over all time points) are used to identify each trench’s central x coordinate and opening end y coordinate. The FOV images are then cropped by a given offset corresponding to trench width and length to generate trench image stacks*. *(c) Example trench image stacks of an acrB-gfp strain. These include the same trench captured simultaneously in three imaging channels: bright field, GFP, and RFP*. *(d) Pre-processing of the fluorescence channels. A bandpass filter is applied to remove the background fluorescence from the feeding lane near trench openings and to reduce pixelation noise. The gradient observed in the GFP intensity profile along the Y-axis is removed as a result*.

To identify trench positions within the individual FOVs, mean intensity profiles in the horizontal (x) and vertical (y) directions are computed (Fig. 3b). Since brightfield images often contain background illumination gradients or diffraction patterns, especially in a microfluidic setting, background subtraction is performed prior to trench identification. Due to the distinctive geometry of this “side trench” mother machine, trench x locations are detected by analyzing the second derivative of the x-intensity profile, which reveals the position of the central and side trench as peaks. For the y-intensity profile, trench opening ends are identified by the strip of diffracted light from the main feeding lane (due to significant depth difference in feeding lane layer and cell trench layer). Using these identified coordinates, individual trenches are extracted and compiled into a structured, multi-dimensional image array, whose dimensions correspond to trench number, time point, imaging channel, trench width, and trench length (Fig. 3c). For a reliable quantification of the fluorescent signals in the next steps, trench images are further processed using a bandpass filter to remove pixelation noise and background gradients coming from the strip of diffracted light from the media channel (Fig. 3d). Resultant filtered images are thereby corrected for long-range fluorescence background signals, and are used in the following steps for extracting single-cell expression analysis.

### Image Processing: Segmentation and Feature Extraction

Machine-learning-based segmentation enables rapid and accurate quantification of single cell morphologies (Cutler et al., 2022; O’Connor et al., 2022). Previously, we have demonstrated the use of Synthetic training data, generated using a virtual microscopy pipeline SyMBac, to train such models for adapting and improving the performance of the machine-learning models in segmenting phase-contrast images. However, segmenting brightfield images is more challenging, due to lower contrast and higher sensitivity to illumination variations and cellular textures. In addition, morphological changes that cells undergo under various conditions such as antibiotic treatments, together with the unique geometry of microfluidic side trenches, make segmentation of treated bacterial cells in brightfield images particularly challenging. While fluorescent segmentation markers improve segmentation accuracy, they compromise temporal resolution due to phototoxicity and photobleaching, reduce available imaging channels for physiological reporters, and impose metabolic burdens on the cells.

To address this challenge, we have developed a two-step training process, to translate the learning from fluorescence images to the corresponding paired brightfield images. Briefly, an Omnipose fluorescence segmentation model is first trained using synthetic image pairs generated using SyMBac (Hardo, 2025b, 2022; Hardo et al., 2022). The resulting model segments sparsely acquired fluorescence images (to minimise phototoxicity and photobleaching) in an experiment.

Segmentation masks generated from these fluorescence images are then combined with simultaneously captured brightfield images (within 200 ms) to generate ground truth pairs for training a brightfield Omnipose model tailored to each experimental setup (Fig. 4a). This brightfield model can subsequently be applied across different experiments performed on the same microfluidic design, without any fluorescent segmentation marker. A similar process has been applied to train a phase contrast segmentation model in our earlier work on tracking bacteriophage infection dynamics on cells (Wedd et al., 2024).

**Figure 4.**
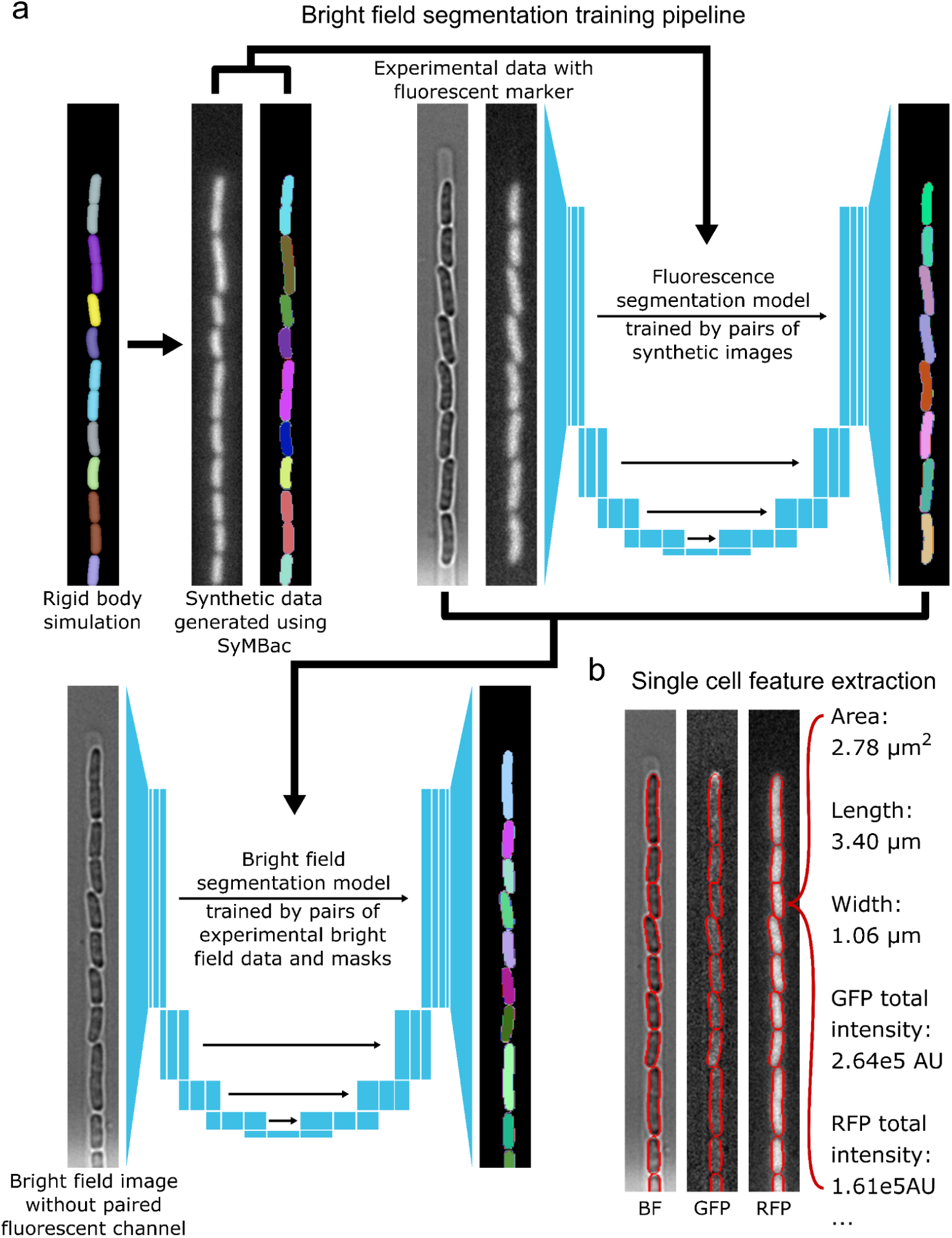
Machine-learning-based segmentation pipeline for brightfield images under microfluidic settings. *(a) Training pipeline for brightfield segmentation. Initially, synthetic fluorescence images with corresponding masks are generated using the SyMBac simulator and renderer. These synthetic images train a fluorescence segmentation model, which is then applied to experimental fluorescence images to produce accurate segmentation masks. These masks, paired with brightfield images captured simultaneously with the fluorescence images, are used to train a brightfield segmentation model specific to the experimental setup. This trained brightfield model can segment brightfield images independently as long as the experimental setup remains consistent, eliminating the need for continuous fluorescence imaging*. *(b) Example of single-cell feature extraction. Segmentation masks from brightfield images enable accurate and frequent measurement of cell morphology including area, length, width, and shape descriptors. The fluorescence intensity for each mask can also be quantified by overlaying the masks on the corresponding GFP and RFP channels*.

These segmentation masks from the individual frames are used for high-throughput, high-temporal-resolution quantification of single-cell morphology (using the scikit-image package for quantifying properties of individual masks through regionprops) and fluorescence intensity across multiple imaging channels from the average and total of pixel intensities within the mask area (Fig. 4b). Accurate and frequent measurements of these properties facilitate precise calculation of growth rates (from changes in mask length over time), generation times (interdivision times on the cell length timeseries), and reporter expression dynamics (from total and average fluorescence timeseries), especially when characterizing bacterial physiological responses under controlled, changing conditions such as sub-lethal drug administration and recovery.

### Lineage Tracking and Reconstruction

To reconstruct bacterial lineages from microfluidic time-lapse data, we developed and utilized a predictive tracking algorithm that simulates the possible states of cell attributes such as size, position, and shape in the next frame (Li, 2024; Wedd et al., 2024). Predictions are guided by exponential cell growth and size regulation models such as adder and sizer, establishing prior probabilities for cell state combinations. Each predicted scenario is matched against observed cell states in the following frame, generating soft-max likelihood probabilities based on the minimum distances achieved when matching the two high-dimensional attribute spaces (Fig. 5a). Positional order constraints within cell trenches are enforced as cells cannot physically switch positions.

**Figure 5.**
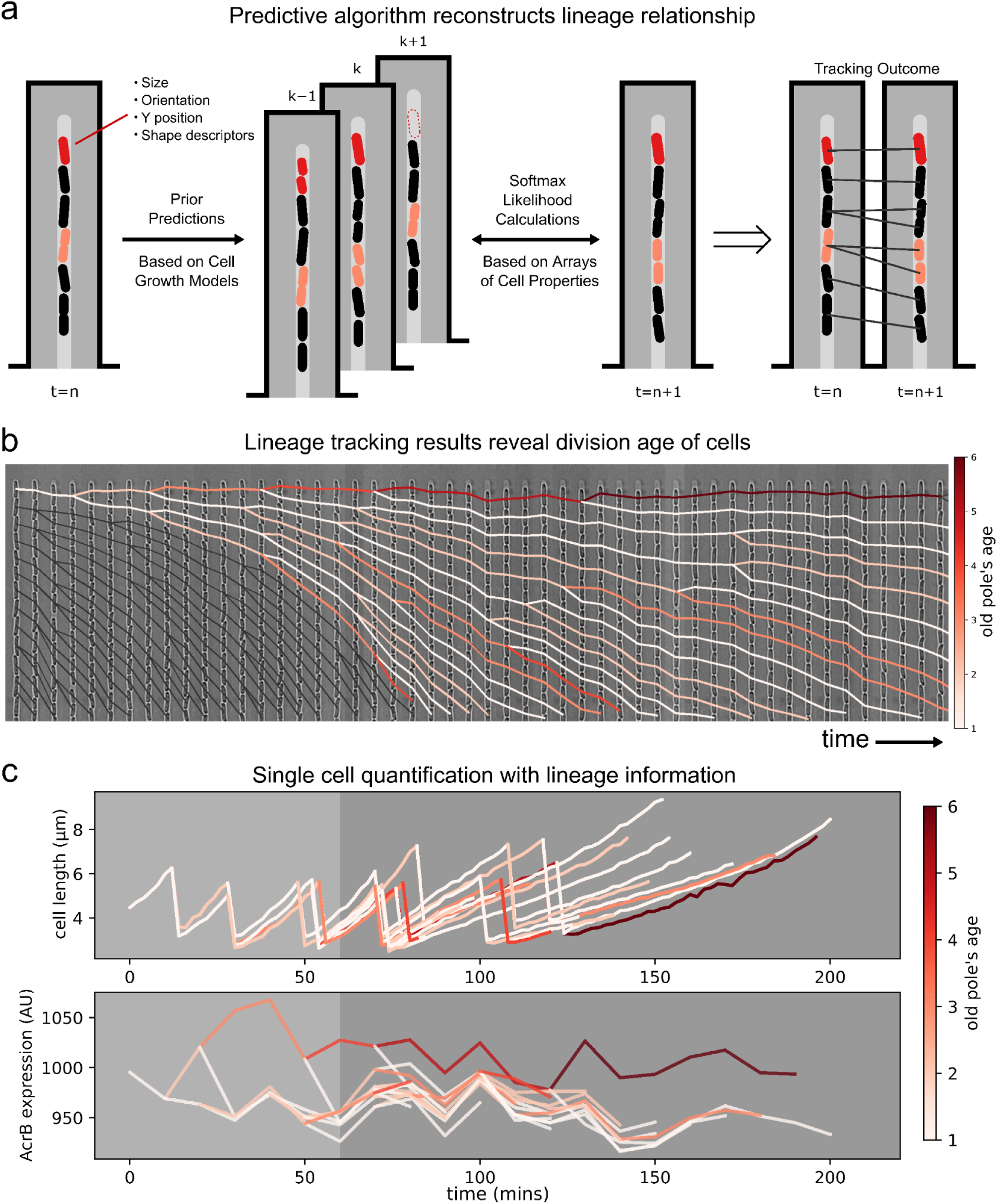
Predictive lineage tracking algorithm for quantifying single-cell response dynamics and their correlation with physiological history. *(a) Schematic of lineage reconstruction algorithm. Cell properties including size, position, and shape are measured in each frame. Their fates in the next frame are simulated based on probabilistic cell growth models with different combinations, generating numerous simulated scenarios. These scenarios are matched to observed cell states in the next frame, returning likelihood probabilities for each match. Final lineage reconstructions are derived from combined prediction and matching scores*. *(b) Example of lineage assignment in time-lapse trench images. Each cell between divisions is color-coded according to the old pole’s age, tracked throughout lineage history. Due to the geometry of the mother machine, the top cell (“mother cell”) has a pole that continuously accumulates pole age with each division*. *(c) Example quantification of lineage dynamics. Cell attributes such as cell size and pump expression are measured and connected between frames, enabling quantification of single-cell dynamics and correlation with history-dependent physiological factors such as pole age*.

Moreover, the algorithm allows the detection of lysis events by permitting skipped cells in the matching process. Final tracking results are obtained by performing Bayesian inference combining prior and likelihood probabilities. Because the number of possible scenarios grows exponentially with the number of cells to track simultaneously and the simulation noise increases proportionally (as positional changes accumulate in one direction), error rates increase with cell number. To minimise such errors, we track a limited number of cells simultaneously, recording their lineage results and looping back. This sequential strategy allows us to track new cells in the subsequent iterations using updated model parameters and previously retained lineages. This significantly improves both algorithmic efficiency and tracking precision.

Using this lineage reconstruction algorithm, we accurately quantify single-cell dynamics including growth rates and efflux pump expression, explicitly considering cell positions within trenches and division ages that drive biased partitioning of efflux pumps. An example timelapse image series is shown in Fig. 5b, where the lineage tracks are overlaid on the brightfield images of cells over time. Individual tracks between two successive division events are colored by the age of the oldest pole of the cell during this period (colormap shown on the right). The cell at the closed end of the trench ages continuously over time, while cells below it follow an oscillating pattern of pole age, going up and down every two positions, due to the nature of the lineage tree resulting from binary fissions. Fig. 5c plots the corresponding growth trajectories (top panel) as cells transition from drug-free condition (light gray) to chloramphenicol stress (dark gray), along with their mean AcrB-GFP expression during an interdivision period (bottom panel), using the same color scheme as Fig. 5b. The quantified traces reveal how cell growth, division frequency, and AcrB expression vary with pole age before and during antibiotic challenge. Noticeably, the mother cell accumulates a large concentration of efflux pumps relative to others. In the following sections, we analyze the relations between a cell’s age, its position, and treatment conditions, to determine how various physiological factors and spatial factors affect the physiological response of individuals.

## Results and discussion

Lineage tracking data reveal the pole age structure of cells within a trench, starting from the first division as they exit from stationary phase, as illustrated in Fig. 6a. Example images from the GFP channel show the distribution of AcrB-GFP efflux pumps within and across cells positioned along a trench. In contrast, the RFP channel displays the distribution of a constitutive control reporter (under the λ phage pR promoter), which exhibits the expected uniform fluorescence across cells and along the cell body. These images are shown alongside a schematic of the expected pole age pattern within a trench. Notably, the GFP signal reveals enhanced AcrB localization at the older poles of certain cells, consistent with polar retention of pumps. These asymmetric distributions are highlighted in the trench image and align with previous findings by Bergmiller et al., who reported biased partitioning of AcrAB-TolC efflux pumps during cell division (Bergmiller et al., 2017). This supports the notion that polar age contributes to the observed heterogeneity in AcrB expression and localization.

**Figure 6.**
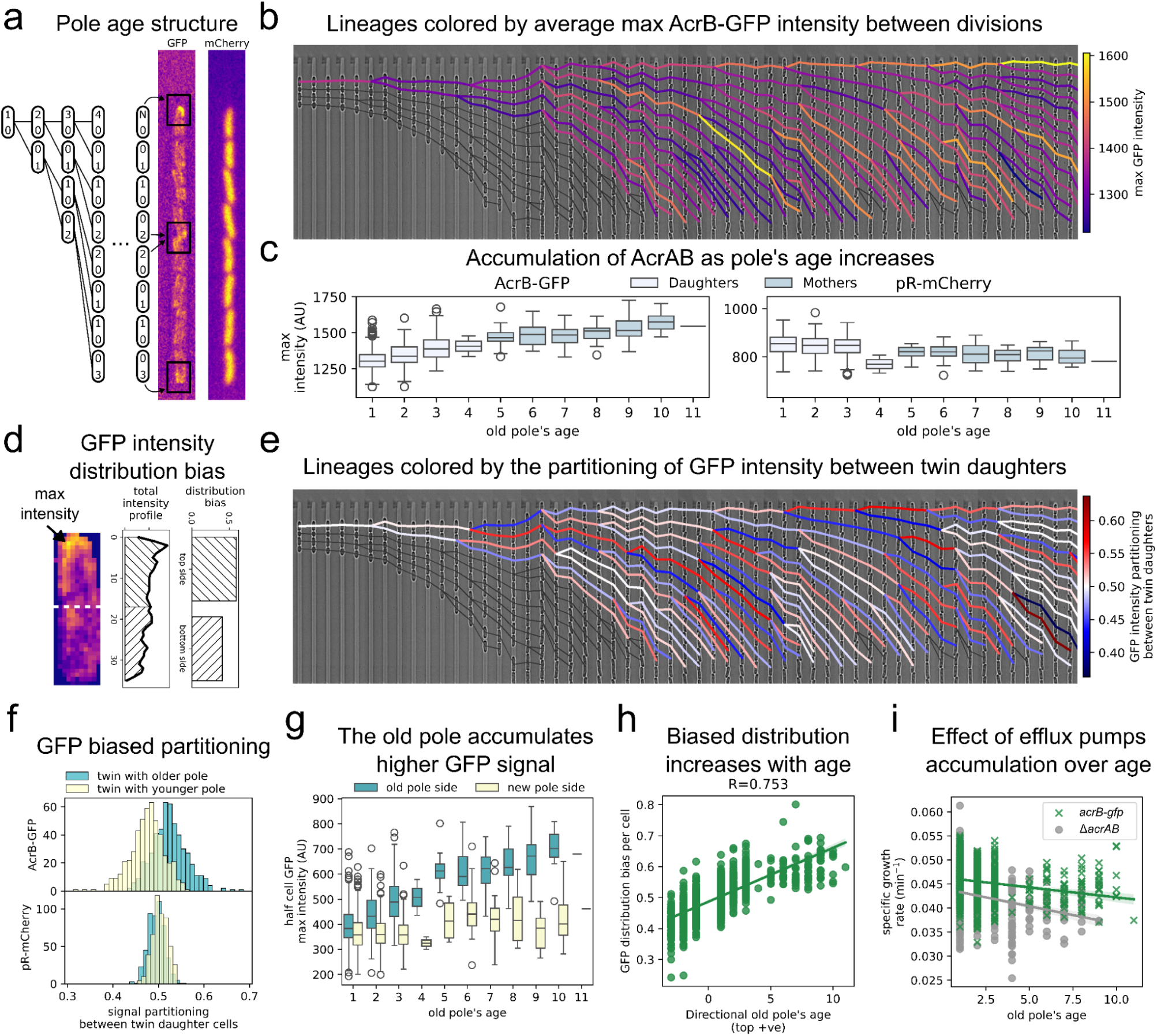
Accumulation and biased distribution AcrAB pump expression leading to biased partitioning similar to pole age structure. *(a) Illustration showing idealized pole age structure in the mother machine under synchronous division condition. Pole age in the mother cell increases with every division, while cells at position 4 and 5 gain pole age 2, and 8th and 9th cell reach pole age 3. Corresponding fluorescence images from the AcrB-GFP and pR-mCherry channels reveal a GFP distribution pattern similar to that of pole age*. *(b) Example lineages from a single acrB-gfp cell, color-coded by their maximum GFP intensity averaged between divisions. The mother cell demonstrates increasing maximum GFP intensity as it transitions from stationary phase into balanced growth*. *(c) Changes in channel maximum intensities with pole age.The AcrB-GFP signals increases with pole age, in contrast to pR-mCherry intensity*. *(d) Quantification of maximum fluorescence intensity and distribution bias. An example single cell AcrB-GFP fluorescence image shown (left), which is extracted by overlaying the segmentation masks on the corresponding GFP channel, with the background subtracted. From this, maximum intensity per cell is measured, and distribution bias is quantified by calculating the fraction of intensities localized in the cell’s top half. In the example shown, 60% of GFP intensity is distributed in the upper half of the cell*. *(e) Example lineages from a single acrB-gfp cell, color-coded by their partitioning of total GFP intensity between twin daughter cells from the same mother. Partitioning is often biased, with up to taking 60% of the intensity while the other twin getting 40%*. *(f) Distributions of partitioning of AcrB-GFP and pR-mCherry. GFP signal preferentially partitions towards daughter cell inheriting older pole than the one with younger pole, contrasting with the more uniform partitioning of pR-mCherry*. *(g) Changes in half cell maximum GFP intensity with pole age. Efflux pump expression accumulates near the old pole*. *(h) Correlation between biased GFP distribution and cell’s pole age. Cells with older poles positioned upwards (positive pole age) exhibit bias values >0.5, confirming that cells inheriting older poles accumulate greater pump levels. This bias becomes more pronounced with increasing pole age, influencing pump partitioning throughout the lineage (p < 0.001)*. *(i) Specific growth rate comparison between the acrB-gfp pump-carrying strain and the ΔacrAB knock-out strain. Cells lacking AcrAB pumps show reduced growth rates, particularly as pole age increases. Accumulation of efflux pumps overall improves growth. (acrB-gfp: R = -0.277, p < 0.001; ΔacrAB: R = -0.231, p < 0.001)*

An example of a tracked lineage within a single trench is shown in Fig. 6b. Even under drug-free conditions, we observe substantial heterogeneity in AcrB-GFP expression dynamics among cells in the same trench. This variability is partially explained by differences in pole age: branches of the lineage tree corresponding to cells with older poles become progressively brighter over time.

Quantitative analysis confirms that AcrB-GFP intensity increases with the age of the old pole (Fig. 6c), while expression from the control reporter (pR-mCherry) remains largely constant across cells, independent of pole age. This age-dependent heterogeneity in AcrB levels arises from the polar localization of the pump in rod-shaped bacteria (Fig. 6d), which leads to asymmetric inheritance during cell division. Following the partitioning pattern across the branches of the lineage tree of an individual trench (Fig. 6e) illustrates this effect: the daughter cell inheriting the older pole, and thus a larger share of polar-localized AcrB-GFP, receives up to ∼60% of the total fluorescent signal (red), while its sibling with the newly formed pole receives ∼40% (blue). This skewed distribution is further validated by population-level histograms of partitioning ratios (Fig. 6f), which show consistent asymmetry in AcrB-GFP inheritance, in contrast to the near-equal distribution of the pR-mCherry control signal. As the population enters balanced growth, an ageing subpopulation of cells emerges with not only higher efflux expression but also increasingly polarized pump distributions (Fig. 6h). The distribution bias between the two new born cells seems to go up with age and then saturate at ∼60%. Because the old pole continuously accumulates more pumps, its fluorescence strengthens over divisions (Fig. 6g), as also seen in the bright yellow branch of the example lineage tree from Fig. 6b.Together, these results highlight how polar localization of efflux pumps and lineage history contribute to physiological heterogeneity even in uniform growth environments.

We next examined the functional implications of asymmetric efflux pump distribution under stress-free, balanced growth conditions. Specifically, we asked whether the presence of AcrAB-TolC pumps imposes a metabolic burden or confers a fitness advantage. To address this, we compared the wild-type strain expressing AcrB-GFP with an isogenic Δ*acrAB* knockout strain. Although growth rates declined with increasing pole age in both strains, the presence of AcrAB conferred a clear benefit: cells expressing the pump maintained higher overall growth rates and exhibited a slower decline in growth with age (Fig. 6i). This suggests that age-dependent accumulation of efflux pumps enhances bacterial fitness even in the absence of external stress. Notably, for cells of the same pole age, those carrying AcrAB pumps grew faster on average than those lacking them. This observation points to a potential role of these pumps in mitigating the accumulation of endogenous toxic metabolites, thereby supporting growth during steady-state proliferation (Teelucksingh et al., 2020).

In Gram-negative bacteria, the AcrAB-TolC system is a major contributor to multidrug resistance. In this study, we focused on the response of *E. coli* to the translation-inhibiting drug chloramphenicol, a known substrate of the AcrAB-TolC efflux pump. Based on growth assays, we estimated the minimum inhibitory concentration (MIC) for chloramphenicol to be ∼12 μg/mL in wild-type cells expressing AcrB. To investigate how cell-to-cell variability in efflux pump expression influences survival and growth under stress, we chose a sublethal concentration of 4 μg/mL. This concentration was sufficient to stress the population while allowing us to explore the protective role of heterogeneous AcrB abundance across individual cells. For comparison, we also examined an isogenic Δ*acrAB* knockout strain, which lacks the ability to actively expel chloramphenicol.

As shown in Fig. 7a, the two strains exhibited markedly different responses to antibiotic treatment. Upon exposure to 4 μg/mL chloramphenicol, nearly all Δ*acrAB* cells ceased growth and failed to recover after drug removal, indicating a critical role of the pump in mediating survival. In contrast, the *acrB-gfp* strain displayed a heterogeneous response: a significant fraction maintained steady-state growth, albeit at a reduced specific growth rate, and then recovered their drug-free exponential growth behavior when the media was switched to antibiotic free media.

**Figure 7.**
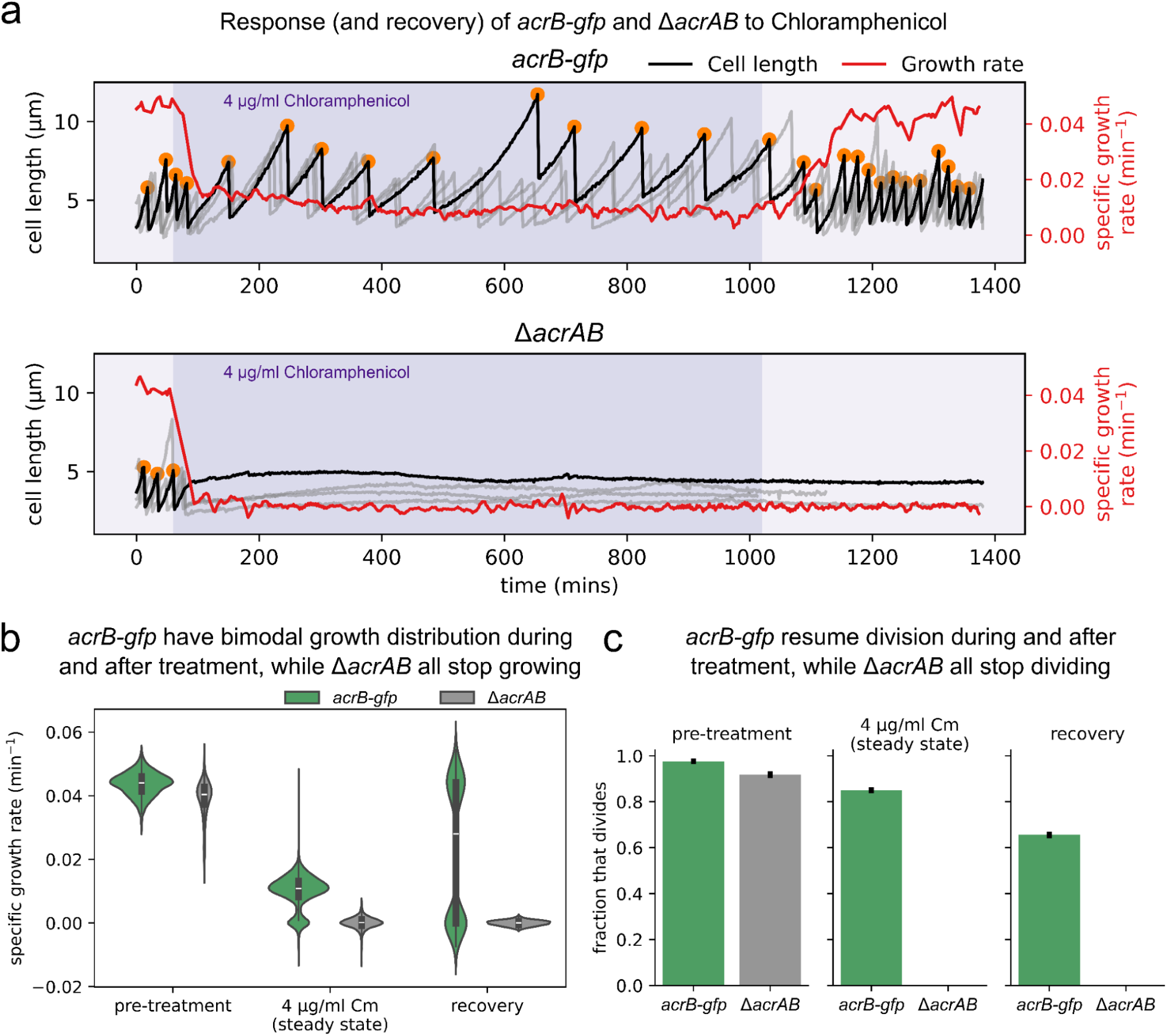
Efflux pump AcrAB enhances bacterial survival under chloramphenicol stress. *(a) Example quantification of mother cell growth trajectories for acrB-gfp and ΔacrAB strain exposed to chloramphenicol and recovery. Bacterial cells in the exponential growth phase (doubling time ∼22 minutes) are introduced to 4 μg/ml chloramphenicol (represented by darker background) at 60 minutes, and are returned to drug-free medium at 1020 minutes. Cell length measurements (at every 2 minutes) enable quantification of division events and specific growth rates (Δlog(cell length)/Δt)*. *(b) Growth response and recovery statistics in acrB-gfp and ΔacrAB cells. All ΔacrAB cells stop growth at steady state during chloramphenicol treatment and fail to recover after antibiotic removal. In contrast, most acrB-gfp cells exhibit reduced but continuous growth at steady state during exposure, with a large fraction returning to pre-treatment growth rates 3 hours after chloramphenicol removal*. *(c) Fraction of dividing cells during and after treatment in acrB-gfp vs. ΔacrAB. In the presence of chloramphenicol, acrB-gfp cells show decreased division activity after reaching steady state, further reduced upon antibiotic removal. Conversely, division halts completely in the ΔacrAB strain, with no recovery observed after drug removal. Error bars represent 95% confidence intervals*.

To ensure unbiased quantification and comparison of this heterogeneity, we focused on mother cells of each genotype, those retained at the closed end of each trench, which are not subject to selective overrepresentation by faster-growing progeny. Growth rates were computed as the change in log-transformed cell length over time (Δlog[length]/Δtime), and division frequencies were recorded (Fig. 7b, c). Prior to treatment, the *acrB-gfp* and *ΔacrAB* strains exhibited growth rates consistent with the patterns observed in Fig. 6i, with Δ*acrAB* cells already showing slightly reduced fitness. During chloramphenicol exposure, the *acrB-gfp* population became bimodal: one subpopulation slowed its growth but continued dividing, while another subset entered a non-growing, arrested state (Fig. 7b). After 17 hours of continuous exposure, the medium was switched back to drug-free growth conditions to simulate the transient nature of antibiotic exposure in vivo, such as antibiotic clearance. Under these recovery conditions, only a subset of the previously arrested *acrB-gfp* cells resumed rapid growth (Fig. 7c). The rest remained arrested or lysed within the 6-hour post-treatment observation window. In contrast, the Δ*acrAB* cells showed no recovery, reinforcing the essential role of AcrB-mediated efflux in surviving even moderate levels of chloramphenicol stress.

To understand how heterogeneity in efflux pump expression impacts antibiotic response at the single-cell level, we analyzed the biological and physical sources of variability within microfluidic trenches using lineage-resolved tracking, spatial position data, and conditional analysis. We focused on three key variables that influence individual cell responses: (1) AcrB-GFP expression level, (2) pole age (a proxy for replicative history), and (3) local antibiotic concentration, which varies along the trench axis (Y position) due to the interplay of diffusion and consumption dynamics (Fig. 8a).

**Figure 8.**
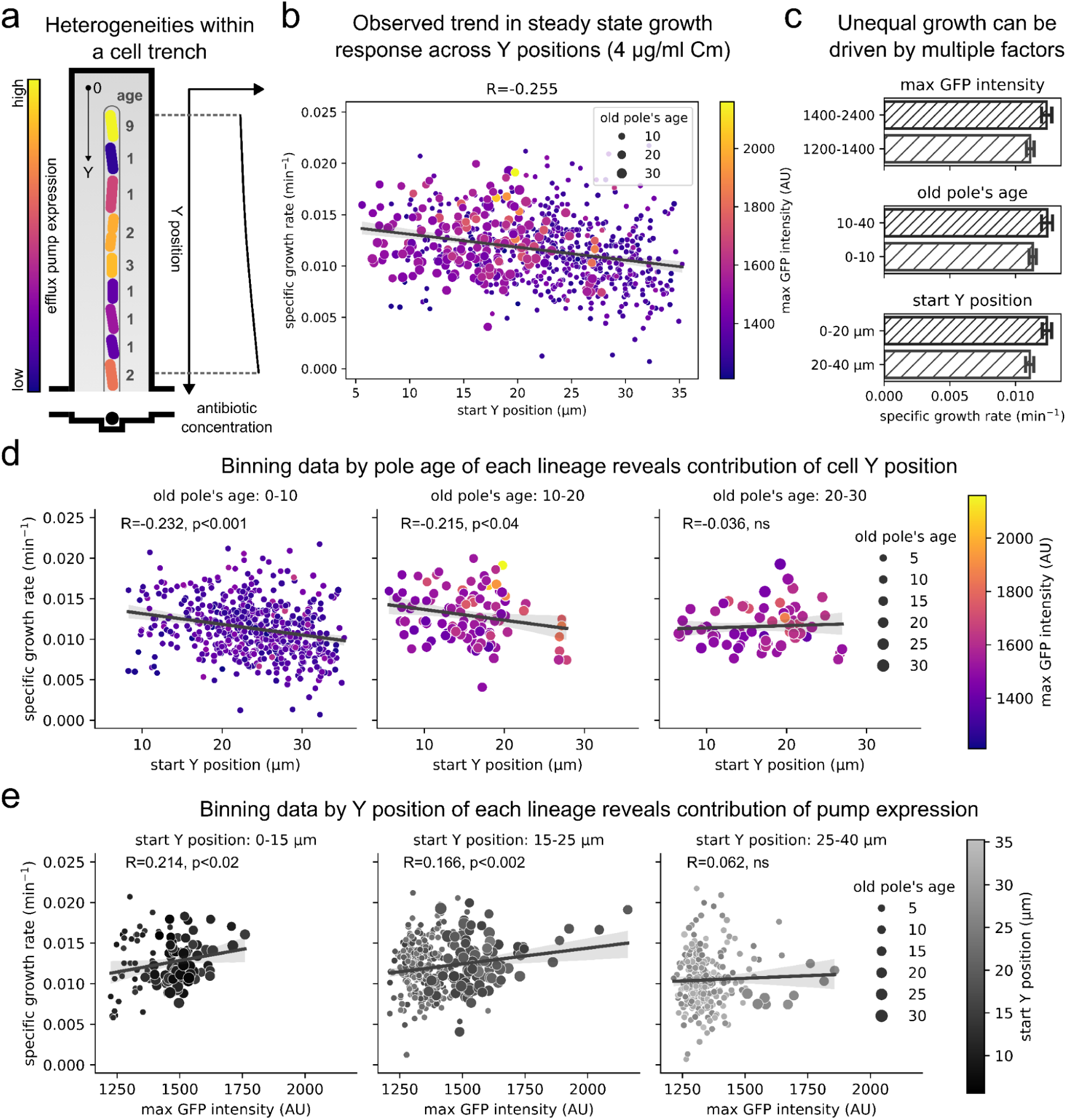
Decoupling factors contributing to heterogeneous bacterial drug responses. *(a) Schematic illustrating the physical and biological heterogeneity within microfluidic trenches. Biased partitioning creates efflux pump heterogeneity similar to pole age structure within cell trenches. Additionally, drug diffusion and cellular titration lead to spatial gradients of antibiotic concentration along the Y direction*. *(b) The scatters show steady state growth rate of acrB-gfp cells in the presence of chloramphenicol, plotted against Y positions recorded at birth of each lineage. Pole age and max GFP expression are represented by scatter size and color respectively. Cells positioned deeper into the trench (lower Y values) have higher growth rates (p < 0.001). However, Y position also associates with pole age and GFP signal, highlighting the need for analyzing conditional dependence of these factors*. *(c) Uneven growth responses in subsets classified by pump max intensity, pole age, and Y position. All three categories have significantly different steady state growth rates in their subsets (p < 0.001, t test). Error bars represent the 95% confidence intervals*. *(d) Effect of Y position on steady state growth rate conditioned by old pole’s age. Conditioned analysis demonstrates that spatial position (likely reflecting antibiotic concentration gradient) impacts growth rates in younger lineages. This effect diminishes in cells with older poles and higher efflux pump expression. (left: R = -0.232, p < 0.001; middle: R = -0.215, p < 0.04; right: ns)*. *(e) Effect of max GFP intensity on steady state growth rate conditioned by start Y position of each lineage. Conditioned analysis reveals a positive correlation between efflux pump expression and steady state growth rate under antibiotic stress. This benefit of pump expression is strongest deeper in trenches, but disappears for cells located closer to the trench opening, where antibiotic exposure is highest. (left: R = 0.214, p < 0.02; middle: R = 0.166, p < 0.002; right: ns)*.

As shown earlier (Fig. 6), cells with older poles, and consequently higher AcrB-GFP levels, tend to reside near the closed (deep) end of the trench. This region is shielded from the direct flow of fresh media, and the delivery of antibiotics becomes limited by diffusion and partially depleted by upstream cells through binding or uptake. Consequently, the concentration of active antibiotics along the trench is non-uniform and less predictable, especially in the lower regions.

To dissect how these factors influence cell behavior during treatment, we visualized the steady-state specific growth rate of AcrB-GFP-expressing cells during antibiotic exposure as a function of their pole age (represented by bubble size), AcrB-GFP level (color-coded), and Y position along the trench (Fig. 8b). The plot reveals that cells near the open end of the trench (higher Y values) tend to have newer poles, lower pump expression, and slower growth rates. In contrast, cells near the closed end are more likely to be older, express more AcrB, and sustain higher growth under antibiotic stress. While growth rate correlates with Y position, suggesting a gradient of antibiotic exposure, it is not immediately clear whether the observed protection is due to increased efflux capacity, spatial shielding from the drug, or replicative age itself (Fig. 8c). These factors are interrelated and must be carefully decoupled to determine causal relationships.

Because lineage tracking provides continuous records of pole age, spatial position, and pump expression for each cell over time, we performed conditional analyses to separate these effects. First, by binning cells based on pole age, we observed that younger cells show a decline in growth rate as they move closer to the trench opening, suggesting that they are more susceptible to spatial variation in antibiotic concentration despite the mitigating effects of the microfluidic design with side trenches (Fig. 8d). In contrast, cells with older poles (e.g., >20 generations) show little to no dependence on Y position, indicating that higher AcrB expression associated with replicative age may buffer them against external drug gradients.

To further isolate the effect of efflux pump expression from spatial variation, we conditioned on Y position and examined the relationship between AcrB-GFP levels and specific growth rate. Among cells positioned away from the trench opening, a positive correlation emerges: cells with older poles and higher AcrB abundance tend to grow faster, while those with newer poles and lower pump levels cluster at lower growth rates (Fig. 8e). Near the opening, where drug concentrations are highest, this correlation becomes weaker, likely because high-efflux, older cells are underrepresented in this region.

Together, these results demonstrate how spatial gradients in antibiotic exposure interact with intrinsic cellular variability, particularly replicative age and efflux pump expression, to shape individual growth outcomes. This layered heterogeneity highlights the need for lineage-resolved, spatially-aware analyses to understand population-level survival and adaptation under antibiotic stress.

## Summary and outlook

Our results highlight how cell-to-cell variability in efflux pump abundance, particularly through asymmetric inheritance of AcrAB-TolC, drives heterogeneous growth, survival, and recovery dynamics under antibiotic exposure. Even at sublethal concentrations of chloramphenicol, we observe a stratified population response: some cells arrest and never recover, others resume growth only after a delay, and a subset continues to divide at reduced rates. The heterogeneous response towards treatment further affects the variable recovery dynamics upon the antibiotic clearance. These phenotypic differences, linked to polar age, efflux expression, and spatial positioning, emphasize that bacterial survival under antibiotic stress is not governed by a single factor but by the dynamic interplay of physiology, inheritance, and microenvironmental exposure. This heterogeneity can serve as a substrate for selection, enabling subsets of the population to persist through treatment and potentially giving rise to future generations with altered resistance profiles, thus bridging short-term antibiotic tolerance and longer-term evolutionary adaptation. These results are broadly consistent with the previous findings from the work by Bergmiller et al. (2017), but provides additional insights from decoupling spatial effects from cellular heterogeneity.

These insights were made possible by a combination of technical innovations: our multilayer side-trench microfluidic device allowed stable long-term tracking of individual cells under realistic, diffusion-limited drug exposure; machine-learning enabled accurate quantifying of cellular phenotypic dynamics, lineage-resolved analysis enabled disentangling inherited traits from environmental effects; and time-lapse fluorescence imaging with controlled treatment profile revealed how efflux pump levels and pole age correlate with growth and survival outcomes. Together, these tools allowed us to observe the functional consequences of efflux heterogeneity with unprecedented resolution.

Looking forward, a key challenge is expanding the range of biological parameters that can be visualized simultaneously in single cells to develop a complete picture of the underlying variables. First, direct visualization of the antibiotic gradient and intracellular accumulation, using fluorescent antibiotic analogs or co-diffusing tracers, would allow quantitative correlation between local drug concentration and cellular response. Second, incorporating reporters for key metabolic states or stress pathways will help uncover how intracellular conditions modulate efflux activity and determine cell fate. Since efflux pump activity is closely tied to energy availability and membrane potential, metabolic reporters are essential to link pump expression with functional capacity. Third, moving beyond AcrAB-TolC to simultaneously track multiple efflux systems or resistance mechanisms (e.g., TolC-dependent and independent pathways) will provide a more complete picture of how bacteria coordinate different survival strategies.

Ultimately, integrating these expanded observables with high-throughput microfluidic imaging and lineage-aware analysis will enable a richer understanding of the molecular logic underlying antibiotic resilience. Such mechanistic insight is critical not only for interpreting single-cell variability but also for informing strategies to design combinatorial treatments that limit persistence and delay resistance evolution.

